# Dokdo sea lion *Zalophus japonicus* genome reveals its evolutionary trajectory before extinction

**DOI:** 10.1101/2024.07.04.602024

**Authors:** Jungeun Kim, Asta Blazyte, Jae-Pil Choi, Changjae Kim, Fedor Sharko, Sungwon Jeon, Eun-Mi Kim, Hawsun Sohn, Jong Hee Lee, Hyun Woo Kim, Mi Hyun Yoo, Kyunglee Lee, Artem Nedoluzhko, Jong Bhak

## Abstract

**Background:** The Dokdo sea lion (*Zalophus japonicus*), commonly referred to as Gangchi in Korea and known as the Japanese sea lion, was endemic to the Northwest Pacific coast before becoming extinct in the 1950s. Little is known about its origins, speciation compared to other *Otariidae* species or how the rapid decline affected the species’ genetic diversity.

**Results:** To raise the Dokdo sea lion from this relative obscurity, we sequenced DNA from 16 *Z. japonicus*’ bone fragments, obtained from Dokdo and Ulleungdo islands in Korea. Our analysis establishes *Z. japonicus* as the earliest diverged species within its genus, significantly redefining its evolutionary relationship with the California (*Z. californianus*) and Galapagos (*Z. wollebaeki*) sea lions. This genome-scale analysis clarifies the phylogeny of *Z. japonicus*, shedding light on its speciation and the evolutionary pathways that shaped its genetic diversity before its extinction. In addition, we discovered, population decline of the *Z. japonicus* started already 1,000 years ago, however, *Z. japonicus* genome maintained a relatively high heterozygosity despite, nearing extinction.

**Conclusions:** Our genome-scale analysis eliminated ambiguity in *Z. japonicus* phylogeny, and shed light on the evolutionary pathways underlying its speciation. This study highlights the importance of the genome-scale analysis for extinct species to understand their complex evolutionary histories and conservation status.

## Background

The Dokdo sea lion – *Zalophus japonicus* (Otariidae: Carnivora), also known as the Japanese sea lion, was a significant species native to East Asia, inhabiting the Dokdo and Ulleungdo islands within Korean territorial waters (Figure 1A). Known locally as Gangchi, this species, along with other Otariidae like the Northern fur seal (*Callorhinus ursinus*) and Steller sea lion (*Eumetopias jubatus*), thrived in the coastal habitats of the northwest Pacific Ocean, spanning from Russia to the coastal waters of Korea and Japan [1]. Despite the absence of dated fossil records for *Z. japonicus*, the discovery of other extant pinniped fossils, such as *E. jubatus*, suggests a long-standing habitation in these regions, potentially since the Pliocene [2]. Historical sources indicate that *Z. japonicus* was hunted for human consumption in Hokkaido since the Jomon period, but it’s widely believed that this subsistence hunting didn’t significantly impact their population dynamics [3]. Over the past two centuries, the *Z. japonicus* population experienced a drastic decline. In the mid-19th century, estimates suggested a healthy population of 30,000 to 50,000 animals, comparable to later counts of Galapagos and California sea lions [3, 4]. However, by the 1950s, their numbers plummeted to just 50-60, leading to their classification as extinct by the International Union for Conservation of Nature (IUCN) in 1990 [3]. This sharp decrease was largely due to extensive hunting for meat, skin, and oil between 1904 and 1925, particularly targeting females and pups [5].

**Fig. 1.**
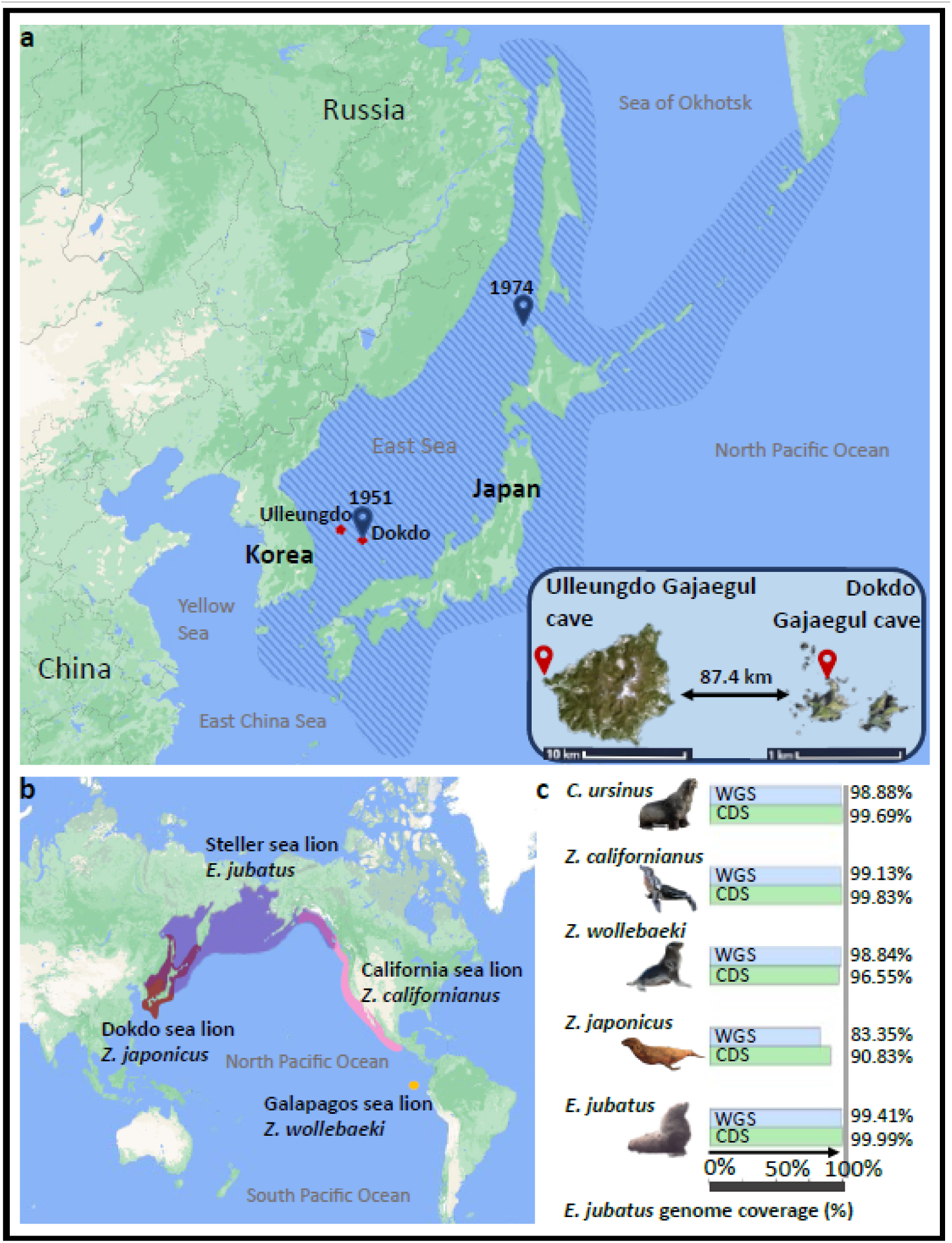
Dokdo sea lion habitat with sampling locations and sequencing (mapping) overview. (a) The map shows the Dokdo sea lion distribution range (striped area), Ulleungdo and Dokdo islands (red dots), and sample sites in Ulleungdo and Dokdo islands (red markers). Samples were excavated from Ulleungdo Gajaegul cave (latitude 37.51° and longitude 130.79°) and Gajaegul cave in western Dokdo (latitude 37.24° and longitude 131.86°). Blue markers mark the two locations where Dokdo sea lions were last seen before being declared extinct and denote the year of last sighting in the location. (b) The species ranges and geographic distribution of *Zalophus* and other Otariidae species relevant in this study. Dark red denotes the species range of the Dokdo sea lion (*Z. japonicus*), pink - species range of California sea lion (*Z. californianus*), and yellow – Galapagos sea lion (*Z. wollebaeki*). Purple semi-transparent area denotes species range of the Steller sea lion (*E. jubatus*), which partially overlaps ranges of Dokdo sea lion and California sea lion. The species distribution range of Northern fur seal (*C. ursinus*) is not shown because it overlaps the range of Steller sea lion almost entirely. (c) Dokdo sea lion (*Z. japonicus*) genome sequencing coverage (breadth of coverage; of protein coding regions – CDS, green, and whole genome – WGS, blue, respectively) comparison with California sea lion (*Z. californianus*) and Northern fur seal (*C. ursinus*), utilizing Steller sea lion (*E. jubatus*) genome as a reference. The maps used for the Otariinae species’ distribution were obtained from https://mapstyle.withgoogle.com. The graphic representations of each species as well as the Ulleungdo and Dokdo islands were created based on royalty-free online images under the Creative Commons (CC) license.

*Z. japonicus* is recognized as a member of the genus *Zalophus*. This genus comprises three distinct species: the western Dokdo sea lion (*Z. japonicus*), the eastern California sea lion (*Z. californianus*), and the *Z. wollebaeki* found from the North Pacific to the Galapagos Archipelago (Figure 1B) [4]. *Z. japonicus* was initially classified as a subspecies of the California sea lion, however, it was reclassified based on unique cranial features [6] and mitochondrial DNA (mtDNA) differences [7]. Evolutionary studies suggest a divergence of merely 2.2 million years between *Z. japonicus* and *Z. californianus*, indicating a close genetic relationship [7]. The specific evolutionary pathways leading to the isolation of these sea lion species remain unclear. Complicating their classification further, pinnipeds exhibit a wide range of individual cranial variations [8] and documented interspecific sexual behaviors [9-12], potentially leading to introgression and morphological diversity. While such behaviors haven’t been directly observed in *Z. japonicus*, the possibility is supported by observed introgression events in other pinniped species [12, 13]. To date, genomic research on *Z. japonicus* has predominantly relied on mitochondrial DNA, which captures only a limited scope of the species’ genomic diversity. Consequently, comprehensive whole-genome sequencing has become essential to fully understand the Dokdo sea lion’s genetic makeup and its phylogenetic relationships within the *Otariidae* [14].

Dokdo Island, a vital habitat for *Z. japonicus* in the 1900s and one of their last observed locations, offers valuable insights. In this study, 16 bone fragments of Dokdo sea lions were unearthed from Dokdo and nearby Ulleungdo islands, situated 87.4 km apart in the East Sea of Korea. We compared the genomes of extinct *Z. japonicus* with those of other sequenced Otariidae species. This analysis not only reveals the genetic diversity and phylogeny of *Z. japonicus* but also sheds light on the evolutionary processes of speciation and introgression in this species for the first time.

## Results

### Dokdo sea lion samples and genomic data

Our study focused on the genomic analysis of Dokdo sea lion using 16 bone samples (Z1, Z3-Z9, Z11-Z18) excavated from Dokdo and Ulleungdo islands (Figure 1a, Additional file 1: Fig. S1, Additional file 1: Fig. S2). These remains, mainly limb and rib bones, underwent DNA extraction and deep DNA sequencing using next-generation sequencing (NGS) methods, despite challenges posed by the small size of the fragments (Additional file 2: Data S1). Detailed information about laboratory techniques used is presented in Supplementary Material. Initial tests on sample Z3 using single-end (SE) and paired-end (PE) NGS revealed higher mapping rates with PE against a *Z. californianus* reference genome, leading us to construct PE NGS libraries for all samples. We generated 8.4 Tb of data with individual mapping rates ranging from 0.1% to 1.3%. Mapping *Z. japonicus* reads to *E. jubatus* genome, the common ancestor of the *Zalophus* genus, covered 83.35% of its genome and 90.83% of protein-coding genes (Figure 1c, Additional file 2: Data S2). This coverage was compared to other Otariidae species, including *Z. californianus, Z. wollebaeki, E. jubatus*, and *C. ursinus*. For ancient DNA bioinformatics analysis, we utilized the PALEOMIX pipeline [15], aligning 43G reads to the *Z. californianus* genome (Additional file 1: Table S1), with an average read length of 140 bp (Additional file 1: Fig. S3). DNA misincorporation levels were low (Additional file 1: Fig. S4), consistent with the species’ recent extinction in the 1950s and 1960s. Additionally, we sequenced modern *Z. californianus* and other pinniped species’ DNA for comparative analysis (Additional file 2: Data S2). This comprehensive genomic approach not only enhances our understanding genome features of *Z. japonicus* but also contributes to the broader knowledge of Otariidae phylogenetics and evolutionary history.

### Congeneric *Zalophus* speciation involved introgression

Dokdo sea lion genetic ancestry and relationship with other *Otariidae* species, is not yet well understood and has not yet been studied outside the morphological classification and mtDNA analysis [7, 16, 17]. To elucidate the genomic composition of *Zalophus* sea lions beyond mtDNA [7, 17], we estimated the gene flow and possible admixture scenarios among *Zalophus* and with other pinnipeds.

Firstly, the *f4*-statistics unsurprisingly show that congeneric *Zalophus* species share the highest genetic affinities among each other compared to the *E. jubatus* (Additional file 1: Table S2). Secondly, *Z. japonicus* exhibits a relatively more distinct genetic makeup compared to its closest species, *Z. californianus* and *Z. wollebaeki* (Fig. 2a-c). In *Z. japonicus*, we could not identify gene flow or introgression from *E. jubatus*, which was observed in both *Z. californianus* and *Z. wollebaeki* (Fig. 2a, b, Additional file 1: Table S3, and Additional file 1: Table S4). Consistently, genetic modeling using qpGraph suggests that *Z. japonicus* diverged early in the evolution of the *Zalophus* genus, with 98% of its genetic components derived from the common ancestor of *Zalophus* and 2% derived from the lineage of *E. jubatus* and *C. ursinus* (Fig. 2c). After the divergence, we observed no additional genetic admixture between *Z. japonicus* and other *Zalophus* species. Third, we suggest that gene flow came from *E. jubatus* and the common ancestor of *Z. californianus* and *Z. wollebaeki* as well as after *Z. japonicus* divergence (Figure 2a, b). *E. jubatus* appeared to share significantly less genetic affinity and gene flow with *Z. japonicus* compared to *Z. californianus* and *Z. wollebaeki* (Figure 2a, b).

**Fig. 2.**
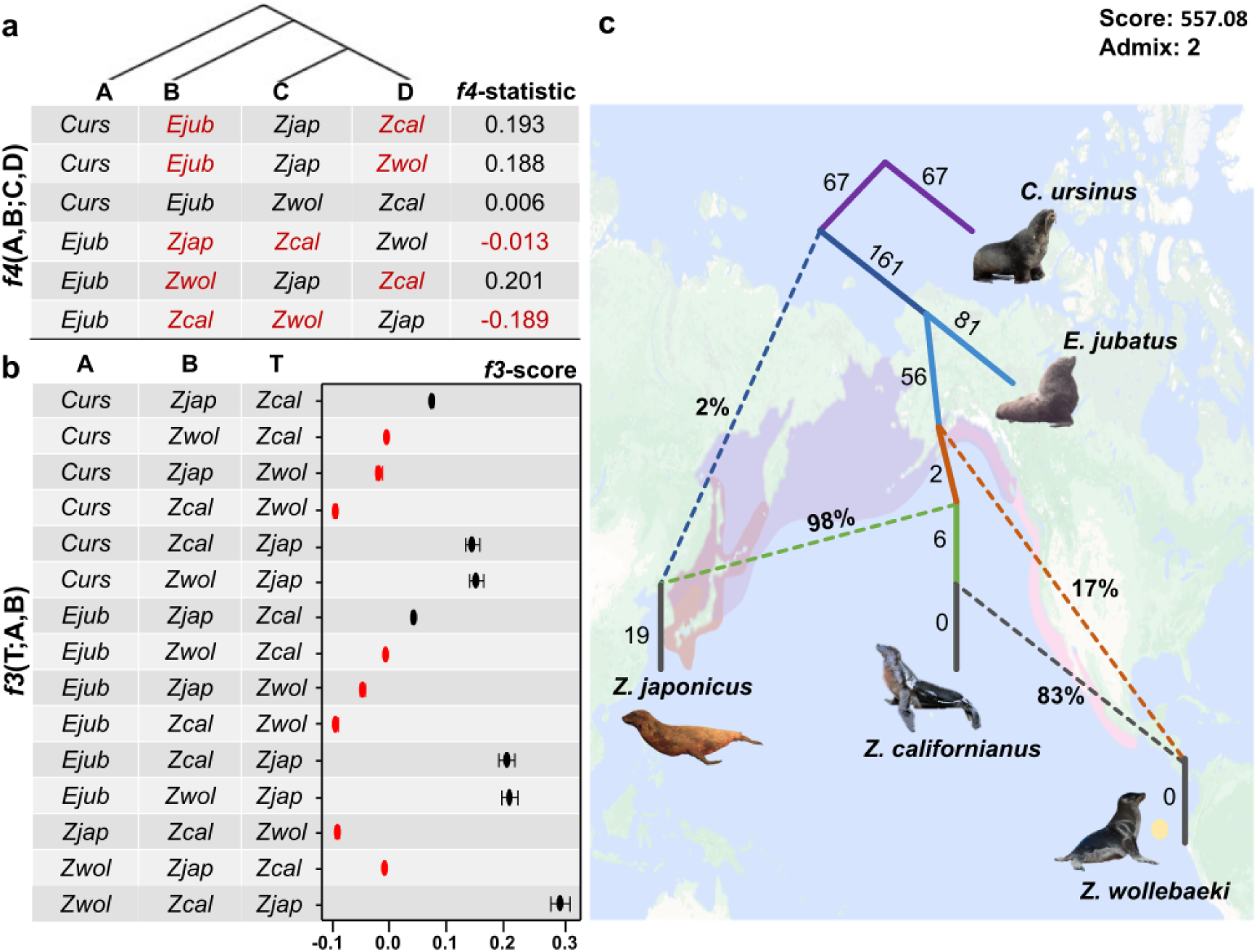
Genetic ancestry of Dokdo sea lion. **(a) Genetic ancestry of *Otariidae* species based on *f4*(A,B;C,D) statistics.** The B genome has a higher genetic affinity with the D genome when the Z-score is > 3. On the other hand, the B genome has higher genetic affinity with the C genome when Z-score is < −3. The red text in the table indicates statistically identical genetic ancestry. The raw data for *f4*-statistics is presented in Additional file 1: Table S3. (b) Admixture *f3* statistics using notation *f3*(T;A,B). In this statistic, negative Z-scores < −3 indicate A and B genomes were admixed in the target genome (T). Instances with negative *f3*-score are presented in red color. The raw data for *f3*-statistics is presented in Table S4. (c) Genetic ancestry of *Zalophus* species based on a qpGraph algorithm [48].

Interestingly, the latter two species not only received more gene flow and derived alleles from *E. jubatus*, but also their levels of genetic affinity to *E. jubatus* were nearly identical (Figure 2a). This provides further support to a scenario wherein *E. jubatus* introgressed into the common ancestor of *Z. californianus* and *Z. wollebaeki* prior to the divergence and isolation of *Z. californianus* and *Z. wollebaeki* as it is quite unlikely that *E. jubatus* introgressed the two modern geographically isolated *Zalophus* species at the same rate (while not sharing any known habitats with *Z. wollebaeki*). Furthermore, following the divergence of *Z. japonicus*, we suggest that more than one major introgression event had occurred between *Z. californianus* and *Z. wollebaeki* (Fig. 2a, b). Lastly, we observed an inconsistent and complex genetic relationship between *Z. japonicus* and other *Zalophus* species, likely due to repeated interspecific and intergeneric introgression events. *f4*-statistics revealed a higher genetic affinity between *Z. japonicus* with *Z. californianus* compared to *Z. wollebaeki* (Fig. 2a). However, admixture *f3*-statistics showed no genetic admixture between *Z. japonicus* and either *C. ursinus* or *E. jubatus* in the *Z. californianus* genome (Fig. 2b). These contradictory finding suggest *Z. californianus* played a significant role in the evolution of the *Zalophus* genus, potentially neutralizing previous admixture effects with *Z. japonicus* through recent genetic interactions with *Z. wollebaeki* (Fig. 2c). In contrast, *Z. wollebaeki* still retains genetic traces of ancestral genetic admixture with *Z. japonicus*. It includes a 17% genetic component shared with all *Zalophus* species (Fig. 2c), indicating a lesser divergence. Additionally, *Z. wollebaeki* did not experience significant genetic exchange with other species (which include introgression from either *C. ursinus* or *E. jubatus*) until its admixture with the common ancestor of *Z. californianus*. These lines of evidence suggest that *Z. japonicus* not only is a unique species with a distinct genetic makeup, that for major part evolved directly from the common ancestor of the *Z. californianus* and *Z. wollebaeki*, but also that *Z. japonicus* may be the earliest diverged species in this genus.

### Complex introgressive speciation of *Zalophus* species explains their phylogenetic ambiguities

Our study for the first time validated *Z. japonicus* phylogeny using 1,581,963 autosomal SNVs (Additional file 2: Data S5) and compared it with whole-mtDNA-based classification (Fig. 3). Both phylogenetic trees presented the same topology with two distinct Otariidae clades and Northern fur seal (*C. ursinus*) as an outgroup: one of Northern pinnipeds composed of *Zalophus* and *Eumetopias* sea lions, and one of Southern pinnipeds composed of *Phocarctos* and *Neophoca* sea lions with *Arctocephalus* fur seals (Fig. 3a, b). In this context, *Z. japonicus* showed almost equal phylogenetic distance to *Z. wollebaeki* and *Z. californianus* (mtDNA: 0.014 and 0.014; WGS: 0.015 and 0.015, respectively) (Additional file 2: Data S6). Even though *Z. japonicus* was similarly related to its congenerics, the genetic distance between the *Z. wollebaeki* and *Z. japonicus* was about 40% greater compared to the distance between the *Z. wollebaeki* and *Z. californianus*. This observation held true regardless of whether the genetic distances were estimated using autosomal SNVs or mtDNA (Additional file 2: Data S6). Our and previously reported [18] intrageneric *Zalophus* phylogeny aligns with *f4*-statistic that showed higher genetic affinity between *Z. wollebaeki* and *Z. californianus* compared to the affinity each of them had with *Z. japonicus*.

**Figure 3.**
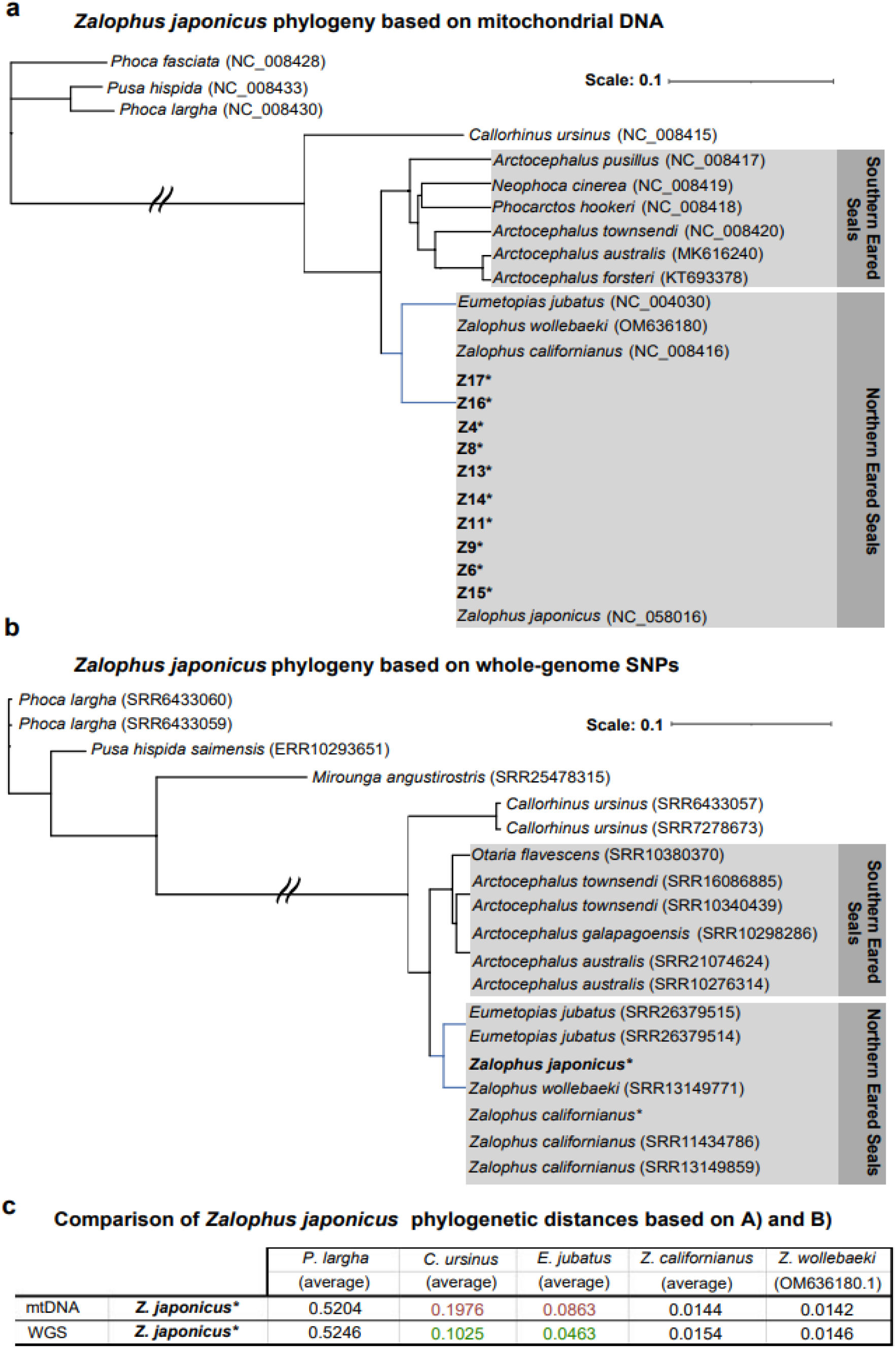
Dokdo sea lion phylogeny. (a) mtDNA-based phylogenetic tree of Otariidae in the context of sea lions and other pinnipeds. The subfamily Otariinae, that consists of *Z. japonicus, Z. wollebaeki*, *Z. californianus*, and *E. jubatus*, is denoted by blue colored branches in the phylogenetic tree. (b) Autosomal SNV-based (WGS) phylogenetic tree of Otariidae in the context of sea lions and other pinnipeds (Additional file 2: Data S5). The subfamily Otariinae, that consists of *Z. japonicus, Z. wollebaeki*, *Z. californianus*, and *E. jubatus*, is denoted by blue colored branches in the phylogenetic tree. (c) Phylogenetic distance comparison between mtDNA and autosomal SNV-based (WGS) phylogeny (Additional file 2: Data S6). Colors mark cases of the genetic distances differing more than 40% between mtDNA and autosomal SNV-based distance matrices. The genetic distances based on mtDNA that are about two-fold higher than those based on autosomal SNVs, are denoted in red. Conversely, the genetic distances based on autosomal SNVs that are about two-fold lower than those based on mtDNA, are denoted in green.

The genetic distances between *Z. japonicus* and its extant genetic donors, *C. ursinus* and *E. jubatus*, showed significant disparities between estimates based on mtDNA and autosomal SNVs (Fig. 3C). The genetic distances based on mtDNA were approximately two times greater than those based on the autosomal SNVs, and reflected in the phylogenetic tree branching (Fig. 3a-c, Additional file 2: Data S5). However, this disparity in genetic distance from *Z. japonicus* did not apply to the already mentioned *Z. japonicus’* congenerics and other phylogenetically distant species, such as those under the genera *Phoca* and *Pusa* (Fig. 3c, Additional file 2: Data S6). These findings imply either a significant divergence of maternal lineages between *Zalophus* and the pair of *C. ursinus* and *E. jubatus* or a genetic legacy of *Z. japonicus* introgression from *C. ursinus* and *E. jubatus*. The introgression scenario obscures phylogenetic relationships, as the relative reduction in autosome-based genetic distance may falsely suggest a more recent common ancestry than it actually is. Nevertheless, both scenarios may be true, and it underscores the multifaceted nature of evolutionary dynamics within the Otariidae, which is also largely consistent with our previously shown genetic ancestry modeling (Fig. 2).

The mtDNA phylogeny of *Z. japonicus* specimens revealed two genetically uniform and nearly indistinguishable mtDNA haplotypes (Additional file 1: Fig. S6). The genetic distances between them were sufficient to identify different mtDNA haplotypes but subtle enough to be possibly derived from the same maternal *Z. japonicus* ancestor (0.0007 vs 0.0005).

### Heterozygosity of extinct Dokdo sea lion

Our subsequent objective was to elucidate the genetic diversity of the *Z. japonicus* samples within the context of population analyses. To accurately estimate heterozygosity (theta) from low depth *Z. japonicus* data, we pooled reads from multiple individuals, enabling genome-wide calculation of heterozygosity across more than 200M loci with a depth of coverage exceeding ten (Additional file 2: Data S6). We obtained moderate heterozygosity value of *Z. japonicus* (0.00101) which was greater than any other *Zalophus* species and several other marine mammal species that are recognized as “Least concern” regarding their vulnerability to extinction (Fig. 4). Moreover, one of *Z. japonicus’* closest living relatives, the non-endangered *Z. californianus*, exhibited about two times lower heterozygosity (0.000491 and 0.000595), suggesting that even a halved estimate for *Z. japonicus* (to compensate for sample merging effect) would not indicate low genetic diversity. Among *Z. californianus*, relatively low heterozygosity value was observed in a potentially inbred individual from a Korean zoo exhibit (0.000491, SRR11434789). This analysis suggests *Z. japonicus’* heterozygosity (0.00101) is not so low as to raise concerns about extinction, and therefore does not indicative severe inbreeding.

**Figure 4.**
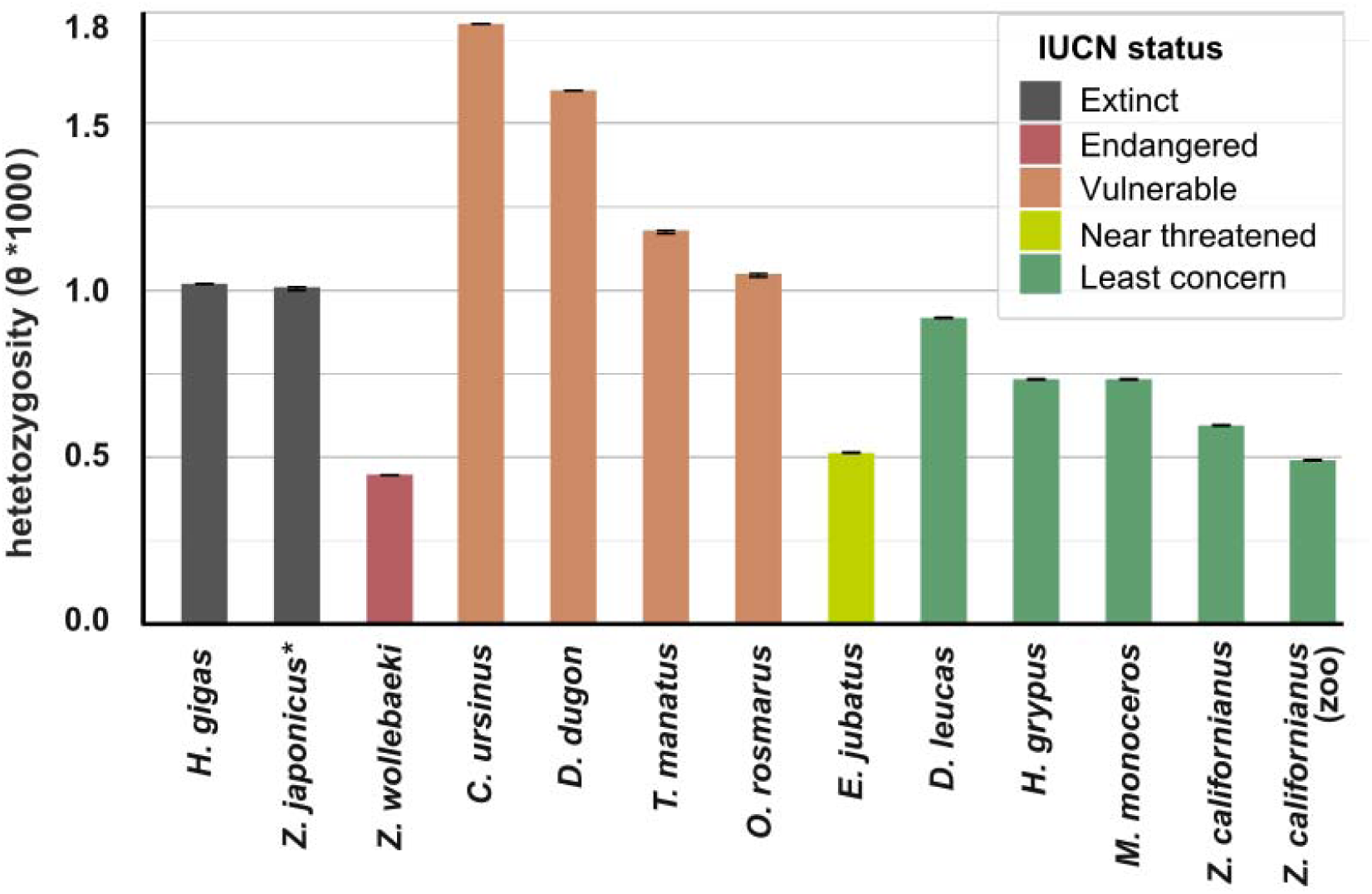
Heterozygosity of Dokdo sea lion in the context of other marine mammals. Logarithm of the average heterozygosity for (from the left to right): Northern fur seal – *C. ursinus*, dugong – *D. dugon*, West Indian manatee – *T. manatus*, alrus – *O. rosmarus*, Steller’s sea cow – *H. gigas*, Dokdo sea lion – *Z. japonicus*, beluga whale – *D. leucas*, grey seal – *H. grypus*, narwhal – *M. monoceros*, California sea lion – *Z. californianus*, Steller sea lion – *E. jubatus*, California sea lion (from zoo) – *Z. californianus*, and Galapagos sea lion – *Z. wollebaeki*. Samples are colored according to present IUCN conservation status.

Interestingly, the heterozygosity levels of *C. ursinus* and *E. jubatus* differ drastically from each other. Despite *C. ursinus* having been extirpated from most of its range over the past 200–800 years due to hunting and environmental factors, its heterozygosity remains at a relatively high level, which corresponds with the historical DNA analysis previously published [19]. In contrast, the heterozygosity estimate for *E. jubatus* is extremely low. We suggest that this fact is related to the nowadays population decline of *E. jubatus*, which began in the 1980s and continues to this day across its distribution range [20, 21].

### Radical population decline of Dokdo sea lion started 1,000 years ago

One crucial determinant of a species’ vulnerability to extinction is its population size, therefore, we inferred the retrospective history of effective population size (*Ne*) changes in *Z. japonicus* and other *Otariidae* species (*Z. californianus*, *Z. wollebaeki*, *E. jubatus* and *C. ursinus*) for comparison, using the pairwise sequentially Markovian coalescent (PSMC) algorithm [22] (Fig. 5). For reliable *Ne* inference over time, we used *Z. japonicus* SNVs identified from genomic loci with a minimum depth of more than ten reads. The population dynamics of *Z. japonicus*, an *Otariidae* species, present a unique pattern compared to its counterparts.

**Figure 5.**
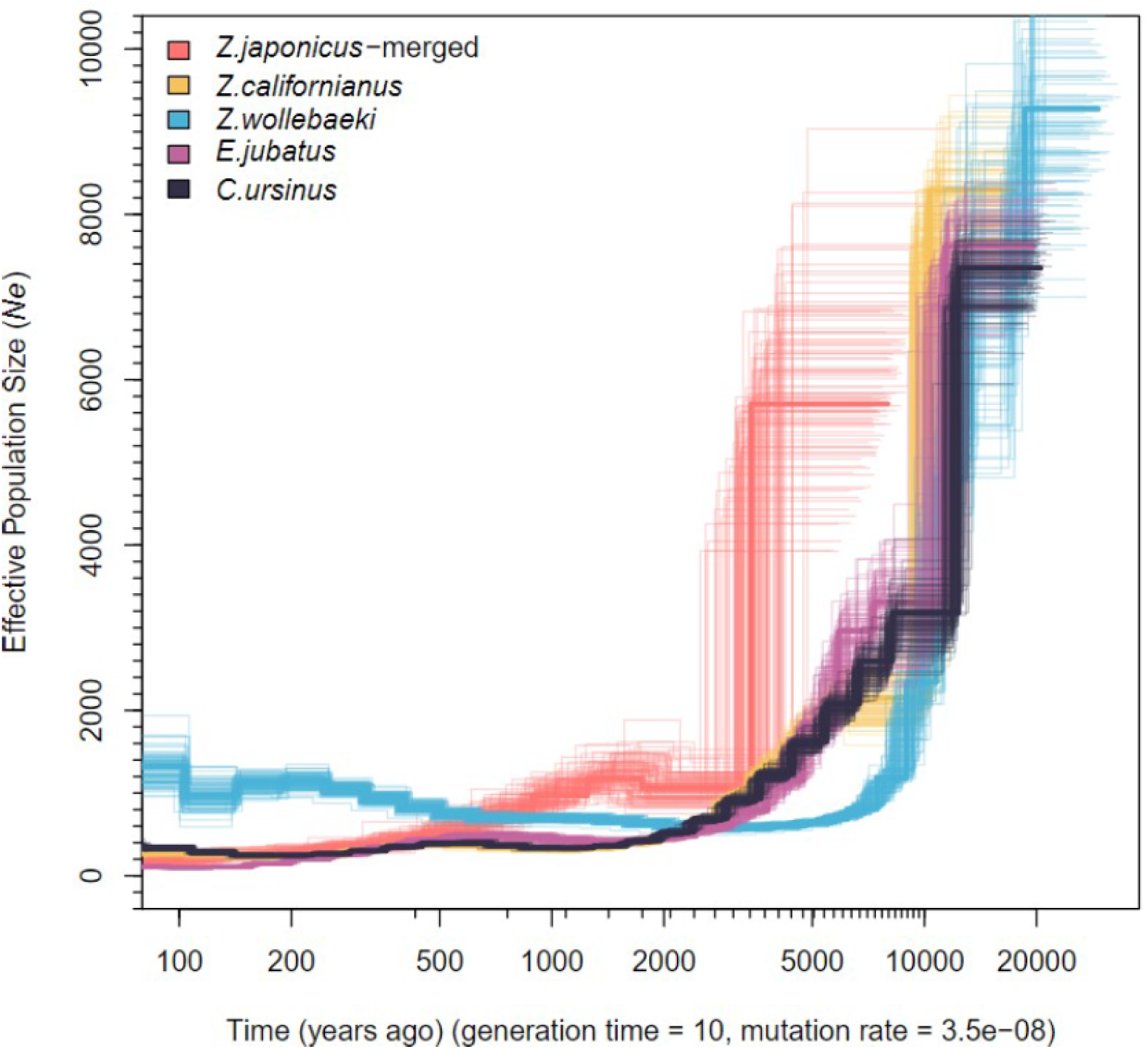
Demographic history of Dokdo sea lion – *Z. japonicus*, California sea lion – *Z. californianus*, Galapagos sea lion – *Z. wollebaeki*, Northern fur seal – *C. ursinus*, Steller sea lion – *E. jubatus*.

While other Otariidae mammals showed a population decline around 10,000 years ago, possibly due to the climatic shifts of the Holocene including warming temperatures and sea level rises, *Z. japonicus* experienced this decline later, around 4,500 years ago. This suggests a different adaptation or response to environmental changes. Notably, the distinct population trajectory of *Z. japonicus* is linked to genetic introgression with *E. jubatus* and *C. ursinus*. We also observed a unique small rebound in population numbers about 1,500 years ago, which was not been observed in the other species. Understanding the interaction between these climatic factors and the population trends of these species is crucial for insights into their evolution and ecological histories. This resurgence was short-lived as a continued decrease in population numbers is apparent ever since.

## Discussion

The development of deep DNA sequencing and increased bioinformatic power and methods has allowed us to understand the basics of evolution better, and to describe traces of genetic introgression and the events that accompanied them, e.g., rapid speciation, multiple ecological radiations, and rapid adaptation to the changing environment [23-25]. Intergeneric fertile hybridization in pinnipeds is well-known fact, which adds an additional layer of complexity analyzing their speciation, phylogeny, and ancestry [12, 26]. Our study sheds light on the evolution extant Otariidae species inhabiting Northern Pacific Ocean and of extinct Dokdo sea lion. Through *f3* and *f4* admixture tests, we describe an introgression from *C. ursinus* and/or *E. jubatus* to *Z. californianus* and *Z. wollebaeki* (Fig. 2a-c). Moreover, we find ancient introgression events between extinct Dokdo sea lion and *E. jubatus/C. ursinus*. While there are no remaining historical or scientific records on *Z. japonicus* hybridization, *Z. californianus* as a species has a rich hybridization history in zoo enclosures with mixed-species pinniped exhibits. On a larger scale, recently, compelling evidence emerged suggesting that smaller wild *E. jubatus* body size found specifically in Oregon population in United States could be attributed to a paternal genetic input from male *Z. californianus* that opportunistically mate with *E. jubatus* females during their seasonal migrations [27]. It is also known that there was a significant overlap not only in the species ranges (Fig. 1b) but also an ecological niche between extinct *Z. japonicus* and extant *C. ursinus*, and *E. jubatus* [3, 4, 28]. We suppose that gene flow between Otariidae species in the north part of the Pacific Ocean and increasing their genetic diversity could have an adaptive effect for these marine mammals in the changing environment for tens of thousands of years and could help them survive at the time of the Pleistocene-Holocene extinction of megafauna.

Unfortunately, increasing human activities over the last few hundred years have led to ecosystem degradation, the destruction of native habitats, and the direct extinction of many animal species. The Dokdo sea lion is one of the species that is the direct victim of human theriocide [3]. Our study confirms the written history, insisting that *Z. japonicus*, a recently extinct iconic species in Korea and Japan, is unique from *Z. californianus* and *Z. wollebaeki* and does not appear to have had a natural evolutionary dead end. Moreover, the demographic history of the Dokdo sea lion has a relatively different trajectory compared with extant Otariidae species. This extinct marine mammal was influenced by a radical decrease in effective population size around 4,500 years ago, while the Northern fur seal, Steller sea lion, California sea lion, and Galapagos sea lion went through genetic bottleneck around 10,000 years ago. Dokdo sea lion is an illustrative example of how human activity can destroy and lead to the verge of extinction of seemingly genetically stable populations of species.

## Materials and methods

### Experimental Design

For our genomic comparison study of the extinct *Z. japonicus*, we used 16 *Z. japonicus* bones (Additional file 1: Fig. S1, Additional file 1: Fig. S2) - three from Gajaegul in Ulleungdo (Gaze Cave, latitude 37.51° and longitude 130.79°) and 13 from Seodo Gajaegul in Dokdo islands (Gaze Cave, latitude 37.24° and longitude 131.86°) (Additional file 1: Table S1). Both sites are named “Gajaegul”, meaning “sea lion cave” in Ulleungdo county dialect. The *Z. japonicus* bones were provided by Cetacean Research Institute of National Institute of Fisheries Science in Republic of Korea. The collection of Dokdo sea lion bones was conducted under the permission granted by the Gyeoungbuk province local government for the collection of protected marine organisms (Permit No. 2019-2). We also collected a *Z. californianus* muscle sample from the Seoul Grand Park, Republic of Korea, obtained during the necropsy process (Permit No. Seoul Grand Park Scientific Research 2020-009). As our study involved extinct animals and cadavers, ethical approval was not required. In addition, we downloaded several 12 pinniped genomes (Additional file 2: Data S5), which used to construct phylogenetic tree, ancestry analysis, and genetic diversity studies.

### DNA extraction and next generation sequencing

To avoid contamination, only endogeneous bone tissue was collected after UV radiation and ethanol treatment. The genomic DNA from the extinct species was extracted using DNeasy Tissue & Blood Kit (Qiagen, Valencia, CA) and the Cetrimonium bromide (CTAB) manual. To generate Illumina NGS data, we constructed PE and SE libraries using the KAPA Hyper Library Preparation Kit (Kapa Biosystems, Woburn, MA, USA) and the Accel NGS 1S plus DNA kit (Swift BioSciences, Washtenaw County, Michigan, USA), respectively, according to manufacturer’s instructions. The Illumina-based NGS sequencing was performed with Illumina NovaSeq 6000 (Illumina, CA, USA) and NextSeq 500 (Illumina, CA, USA). For MGI-Seq, we constructed paired-end libraries with MGIEasy DNA Library Prep Kit (MGI, Shenzhen, China) and sequenced them on the DNBSEQ-T7 sequencing platform.

### Dokdo sea lion mitochondrial genome assembly

Upon assessing the read quality with Trimmomatic (ver. 0.39) [29], we extracted mitochondrial DNA (mtDNA) reads by mapping all the short DNA reads to the mito-genome of *Z. californianus* (Acc. NC_006416). We assembled the mitochondrial genome (mito-genome) of the *Z. japonicus* (Z6) sample using NOVOPlasty (ver. 4.2) [30]. Minor gaps of the mito-genome assembly were filled in by conducting Sanger sequencing (Additional file 1: Table S5, Additional file 1: Table S6) followed by assembly using Cap3 program (updated on December 21, 2007) [31]. We predicted and annotated *Z. japonicus’* mito-genome using the MITOS program (http://mitos.bioinf.uni-leipzig.de/index.py) (Additional file 1: Fig S5, Additional file 2: Data S3) [32]. Complete mito-genomes were aligned with Mummer (ver. 4.0.0rc1) (Additional file 1: Fig. S5) [33] and Dendroscope (ver. 3.5.10) [34]. We additionally obtained ten mito-genome consensus sequences by aligning our samples’s NGS reads to Z6 deep-sequenced mito-genome (Additional file 2: Data S4). The consensus sequences were obtained from high quality SNV data (mapping quality > 30, genotype quality > 20, and coverage > 10) by implementing samtools consensus utility (ver. 1.9) [35]. Five samples (Z1, Z5, Z7, Z8, and Z12) were excluded in the process due to significantly lower (insufficient) amount of NGS reads (Additional file 2: Data S4). We then constructed the phylogenetic tree of the ten *Z. japonicus* mito-genomes along with other closely related species (Fig. 3a, Additional file 1: Fig. S6 and Additional file 2: Data S6). We aligned CDS sequences using muscle program (ver. 3.8.31) [36] and constructed the phylogenetic tree using phyML (ver. 3.1) with default parameters (ver. 3.1) [37].

### Phylogeny, admixture and genomic composition analyses of Dokdo sea lion and other pinnipeds

To construct phylogenetic tree and analyze ancestry, we aligned reads to an outgroup species that is the most distantly related to *Z. japonicus*, namely, the walrus, *Odobenus rosmarus* (acc. ANOP00000000) [38]. Specifically for ancient DNA samples, we applied PALEOMIX pipeline [15] by mapping all *Z. japonicus* reads to the *O. rosmarus* genome [38]. We estimated deamination patterns using mapDamage (ver. 2.0) [39]. For the modern mammal genomes, their NGS reads were aligned to the same reference genome using bwa mem (ver. 0.7.17) [40] after filtering out low quality reads using Trimmomatic with Quality < 30 and read length < 70 (ver. 0.39) [29]. We then utilized Picard [41] (ver. 2.27.5) to eliminate PCR duplicates and employed the GATK (ver. 4.1.3.0) for variant calling [42]. We constructed consensus sequence using the vcf2phylip [43] utilizing only genomic loci in the *Z. japonicus* bam files with read depth (DP) larger than five. We then constructed phylogenetic tree using the PhyML (ver. 3.1) [37] as in mitochondrial genome.

The *f3-* and *f4-*statistics (table S2-4) were conducted using a ADMIXTOOLS (ver. 2.0) algorithm [44]. An admixture graph was constructed with qpGraph model [44] with admix=2. The qpGraph [44] was automatically optimized for genetic admixture of our admixture model.

### Estimation of the effective population size of Dokdo sea lion

We used PSMC algorithm [22] to estimate the effective population size of Dokdo sea lion in the last 20,000 years. For higher mapping rates, for this analysis, we aligned reads from all ancient and extant *Zalophus* and *E. jubatus* species to the *E. jubatus* reference genome (acc. GCA_004028035.1, ver. ASM402803v1), which shares more recent common ancestry with these species compared to the previously used walrus. Using the PALEOMIX pipeline [15] we aligned all *Z. japonicus* reads to the *E. jubatus* reference and selected only high confidence SNVs, with more than 10x coverage for the PSMC analysis. The stringent data pre-filtering aimed to reduce biases stemming from possible over-representation of heterozygous sites in the ancient DNA. This was conducted as a necessary step, because PSMC [22] infers the *Ne* changes over time using the density of heterozygous sites throughout the diploid genome of a single individual. For the PSMC analysis of modern genomes, we aligned reads with bwa mem (ver. 0.7.17) [40] after trimming low quality reads using Trimmomatic (ver. 0.39) [29]. We proceeded by utilized Picard (ver. 2.27.5) to eliminate PCR duplicates (https://broadinstitute.github.io/picard/) and employed the GATK (ver. 4.1.3.0) for variant calling [42]. We applied a generation time of 10 [45] and a mutation rate of 0.27 * 10^-8^ [46].

### Calculation of the heterozygosity of Dokdo sea lion genomes

To accurately identify the heterozygous regions in the *Z. japonicus* genome, we calculated the distribution of heterozygous positions across the genomic loci from a bam file, wherein Z. japonicus reads were mapped to the most closely related reference genome, *Z. californianus* (Acc. GCF_00976235.2) using the PALEOMIX pipeline [15]. We called SNVs utilizing samtools mpileup [35] with a minimum base quality of 20 (-Q 20) and mapping quality 20 (-q 20). We included genomic loci with a DP of 10 or greater. We cleaned all reads from the modern mammal genomes using the Trimmomatic (ver. 0.39) [29] and mapped them using the bwa mem (ver. 0.7.17) [40] to the most closely related reference genomes (Additional file 2: Data S7). After removing the PCR duplicates using Picard (ver. 2.27.5), we applied the same mpileup criteria with *Z. japonicus*. This way, we implemented mlRho (ver. 2.9) [47] to estimate the heterozygosity only on the high-quality loci with high quality variants.

### *f*-statistics analyses for Dokdo sea lion and other *Otariidae* species

For ancestry analysis, we used *f4*-statistics and admixture *f3*-statistics. The admixture *f3*- and *f4*-statistics were conducted using a ADMIXTOOLS (ver. 2.0) algorithm [44]. All formulars employed are detailed in Additional file 2: Data S5-S7. Our criterion for significance was set at an absolute Z-score less than three.

## Declarations

### Ethics approval and consent to participate

The collection of Dokdo sea lion bones was conducted under the permission granted by the Gyeoungbuk province local government for the collection of protected marine organisms (Permit No. 2019-2). We also collected a *Z. californianus* muscle sample from the Seoul Grand Park, Republic of Korea, obtained during the necropsy process (Permit No. Seoul Grand Park Scientific Research 2020-009). As our study involved extinct animals and cadavers, ethical approval was not required.

### Consent for publication

Not applicable

### Availability of data and materials

All data are publicly available for scientific research. Sequencing data have been deposited in the NCBI SRA with accession numbers PRJNA982545. All data are available in the main text or the supplementary materials.

### Competing interests

C.K. and S.J. are employees and J.B. is the CEO of Clinomics Inc. Other authors declare that they have no competing interests.

### Funding

National Institute of Fisheries Science, Ministry of Ocean and Fisheries, Korea (R2020024, R2021030, R2022033, R2024004). Promotion of Innovative Business for Regulation-Free Special Zones funded by the Ministry of SMEs and Startups (MSS, Korea) (grant number [P0016195, P0016193] (1425156792, 1425157301) (2.220035.01, 2.220036.01)). Ulsan City Research Fund (1.200047.01). Fedor Sharko and Artem Nedoluzhko were supported by ARCTIC SIRENIA RESEARCH FOUNDATION.

### Author’s contributions

Conceptualization: J. K., J. C., A. B, S. J and F. S., A. N.; Methodology: C. K., J. K, F. S; Investigation: E. K, H.-W. K., M. Y., J.-H. L, K. L., and H. S.; Visualization: A. B, J. C., J. K.; Supervision: J. K, A. B; Writing—original draft: J. K., J. C., A. B; Writing—review & editing: A. N, J. B

## Acknowledgements

Not applicable

## Supplementary Information

### Additional file 1

**Fig. S1.** Excavation of Dokdo sea lion bones

**Fig. S2.** Dokdo sea lion bones used in this study

**Fig. S3.** Read length distribution mapped to California sea lion (*Z. californianus*) genome

**Fig. S4.** Postmortem DNA damage pattern in the DNA-libraries of Dokdo sea lion generated by PALEOMIX pipeline

**Fig S5.** Assembly and annotation of the complete mitochondrial genome of Dokdo sea lion using the Z6 individual specimen

**Fig. S6.** Phylogenetic tree of the mitogenomes of Dokdo sea lion and related species

**Fig. S7.** Comparison of mitochondrial genomes between *Z. japonicus* and *Z. californianus* using mummer (ver. 4.0.0rc1)

**Table S1.** Statistics of the PALEOMIX pipeline mapping to California sea lion (*Z. californianus*) reference genome

**Table S2.** Gene flow between Steller sea lion (*E. jubatus*) and *Zalophus* species with a form of *f4*(A,B;C,D).

**Table S3.** *f4*-statistics of *Otariidae* species with a form of f4(A,B;C,D).

**Table S4.** Genetic admixture of *Zalophus* species compared to their related species.

**Table S5.** Primers to filling the mito-genome gap of Dokdo sea lion (Z6 individual) and melting temperature (Tm) used

**Table S6.** Sanger sequencing reads to fill mito-genome gap of Z6 individual

### Additional file 2

**Data S1**. Statistics of DNA sequencing and mapping statistics to California sea lion reference genome.

**Data S2**. Number (No) of mapped read to Steller sea lion reference genome and coverage statistics.

**Data S3.** Annotation of the mitochondrial genome of Dokdo sea lion.

**Data S4.** Coverage of the mitochondrial genomes based on reads with mapping quality > 30, genotype quality > 20, and coverage >10.

**Data S5.** Mammal genome datasets used for the ancestry analysis.

**Data S6.** Genetic distance matrix for mtDNA (A) and nuclear genome (B).

**Data S7.** Average genome-wide autosomal heterozygosity values.

### Rights and permissions

Open Access This article is licensed under a Creative Commons Attribution 4.0 International License, which permits use, sharing, adaptation, distribution and reproduction in any medium or format, as long as you give appropriate credit to the original author(s) and the source, provide a link to the Creative Commons licence, and indicate if changes were made. The images or other third party material in this article are included in the article’s Creative Commons licence, unless indicated otherwise in a credit line to the material. If material is not included in the article’s Creative Commons licence and your intended use is not permitted by statutory regulation or exceeds the permitted use, you will need to obtain permission directly from the copyright holder. To view a copy of this licence, visit http://creativecommons.org/licenses/by/4.0/. The Creative Commons Public Domain Dedication waiver (http://creativecommons.org/publicdomain/zero/1.0/) applies to the data made available in this article, unless otherwise stated in a credit line to the data.

