## Additional file 1 - supplementary figures for "Dokdo sea lion *Zalophus japonicus* genome reveals its evolutionary trajectory before extinction"

Jungeun Kim ^1†^ *et al.*

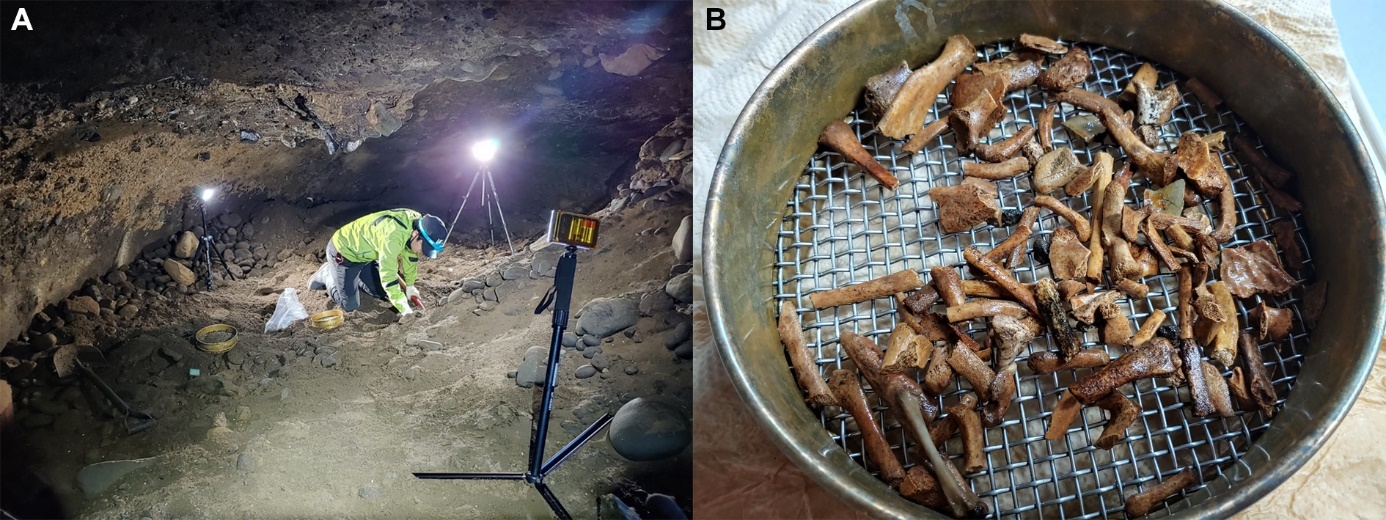

**Fig. S1. Excavation of Dokdo sea lion bones**

(A) *Z. japonicus* bones excavation site (Gazegul cave in western Dokdo Island (latitude 37.24° and longitude 131.86°), (B) *Z. japonicus* bones excavated.

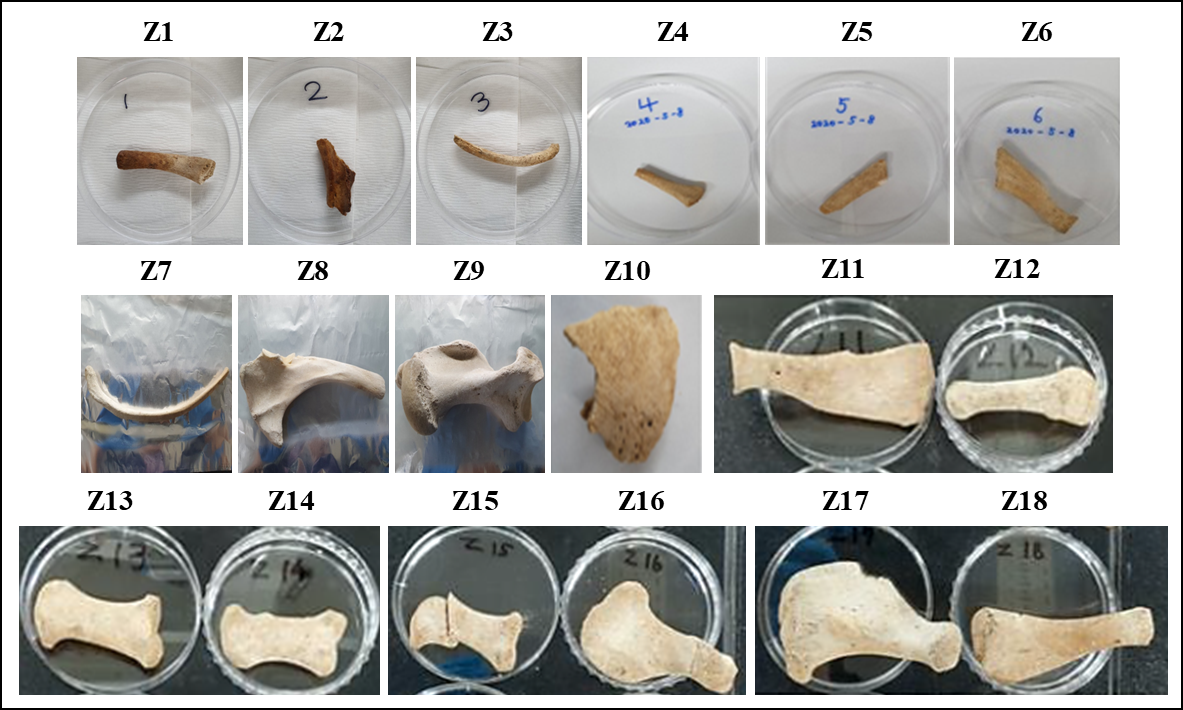

**Fig. S2. Dokdo sea lion bones used in this study.**

Dokdo sea lion samples: Z1~Z6, and Z10~Z18, Ulleungdo samples: Z7~Z9 (table S1).

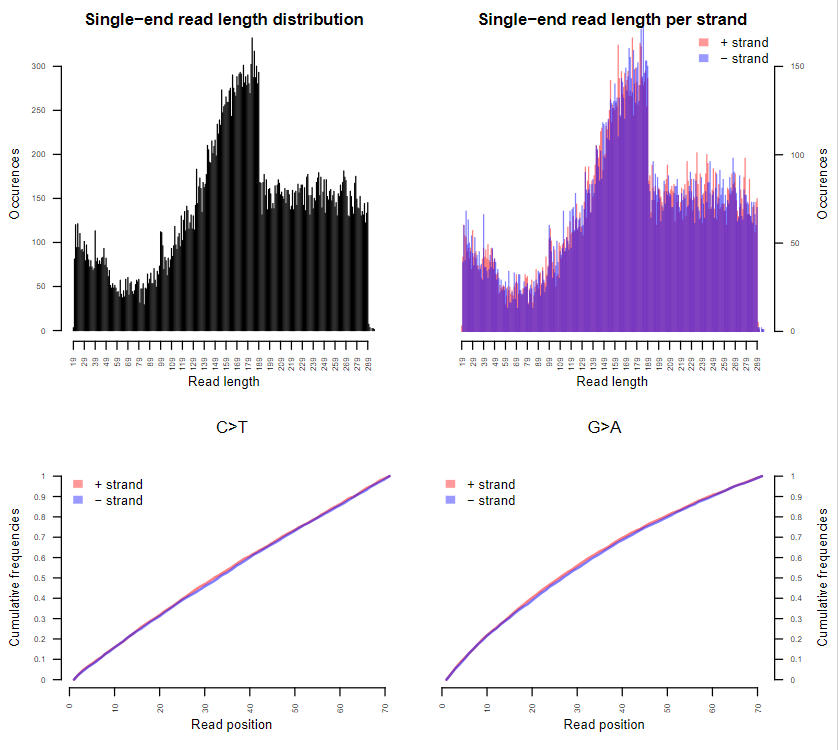

**Fig. S3. Read length distribution mapped to California sea lion (*Z. californianus*) genome**

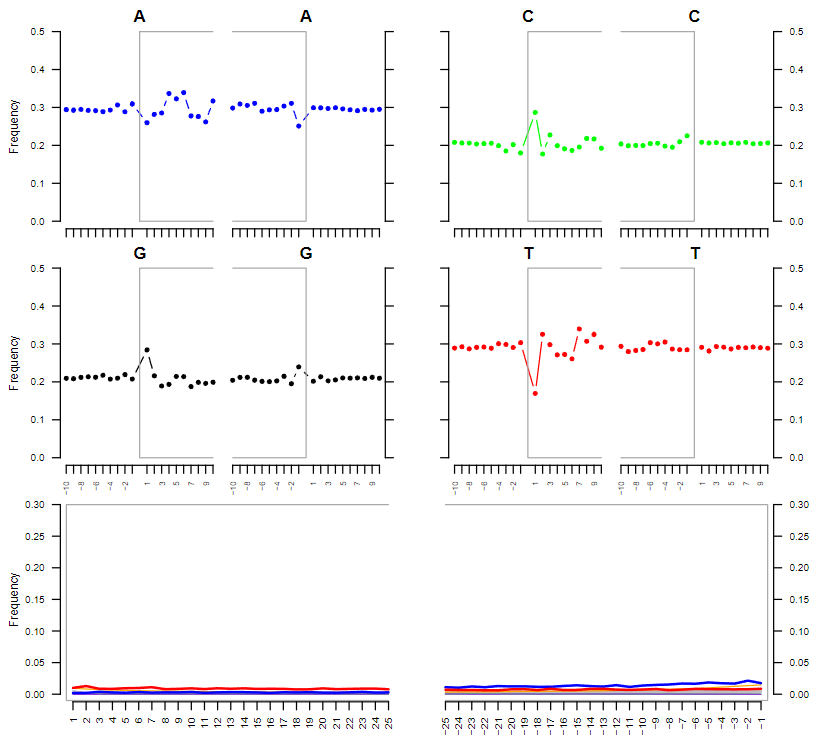

**Fig. S4. Postmortem DNA damage pattern in the DNA-libraries of Dokdo sea lion generated by PALEOMIX pipeline present C to T (and complementary G to A) misincorporations at the 5’ and 3’ termini of the terminal 25 nucleotides. The Y-axis shows frequency of misincorporations. The X-axis shows last 25 nucleotides of DNA libraries**

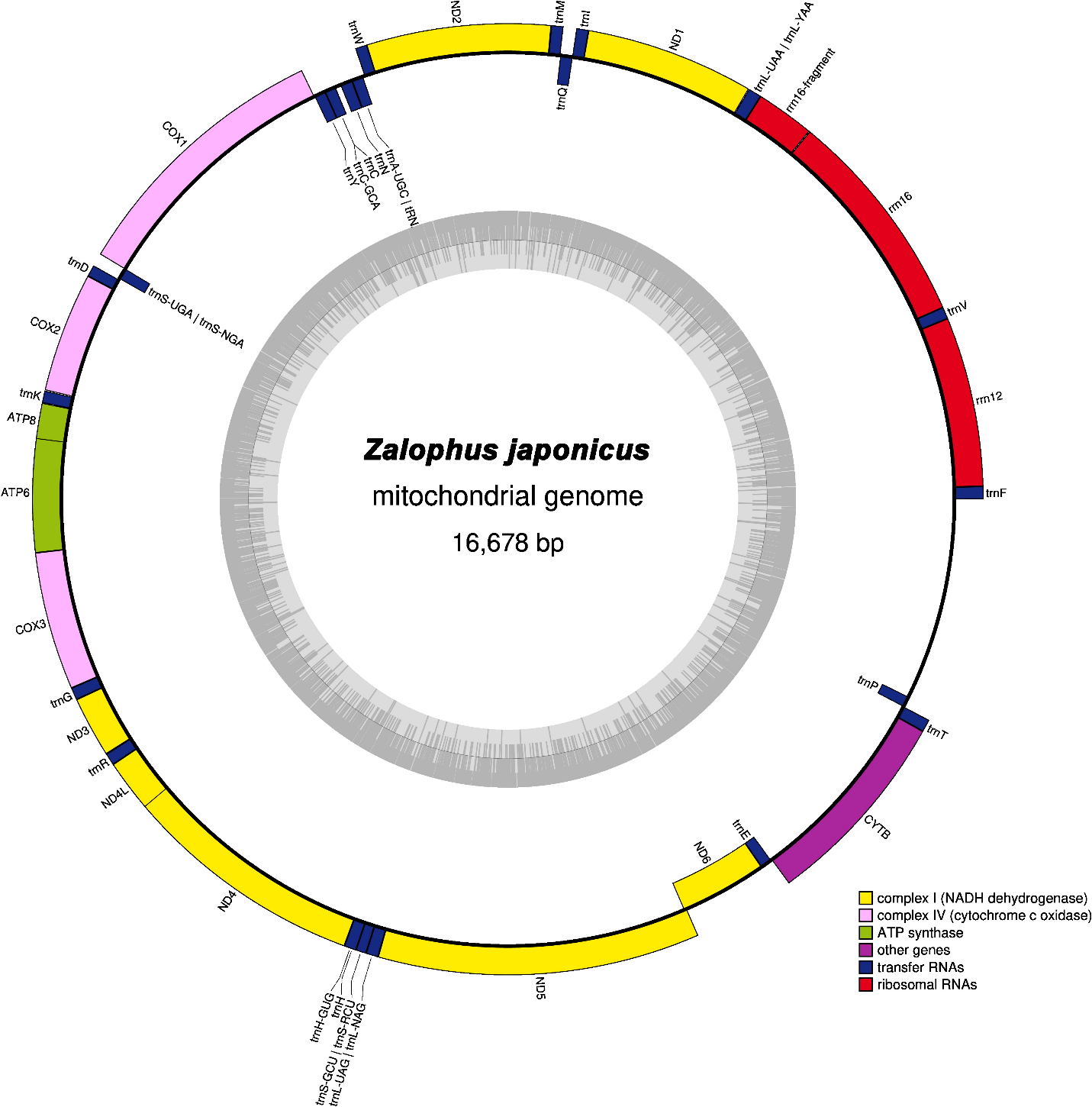
 **Fig. S5. Assembly and annotation of the complete mitochondrial genome of Dokdo sea lion using the Z6 individual specimen**

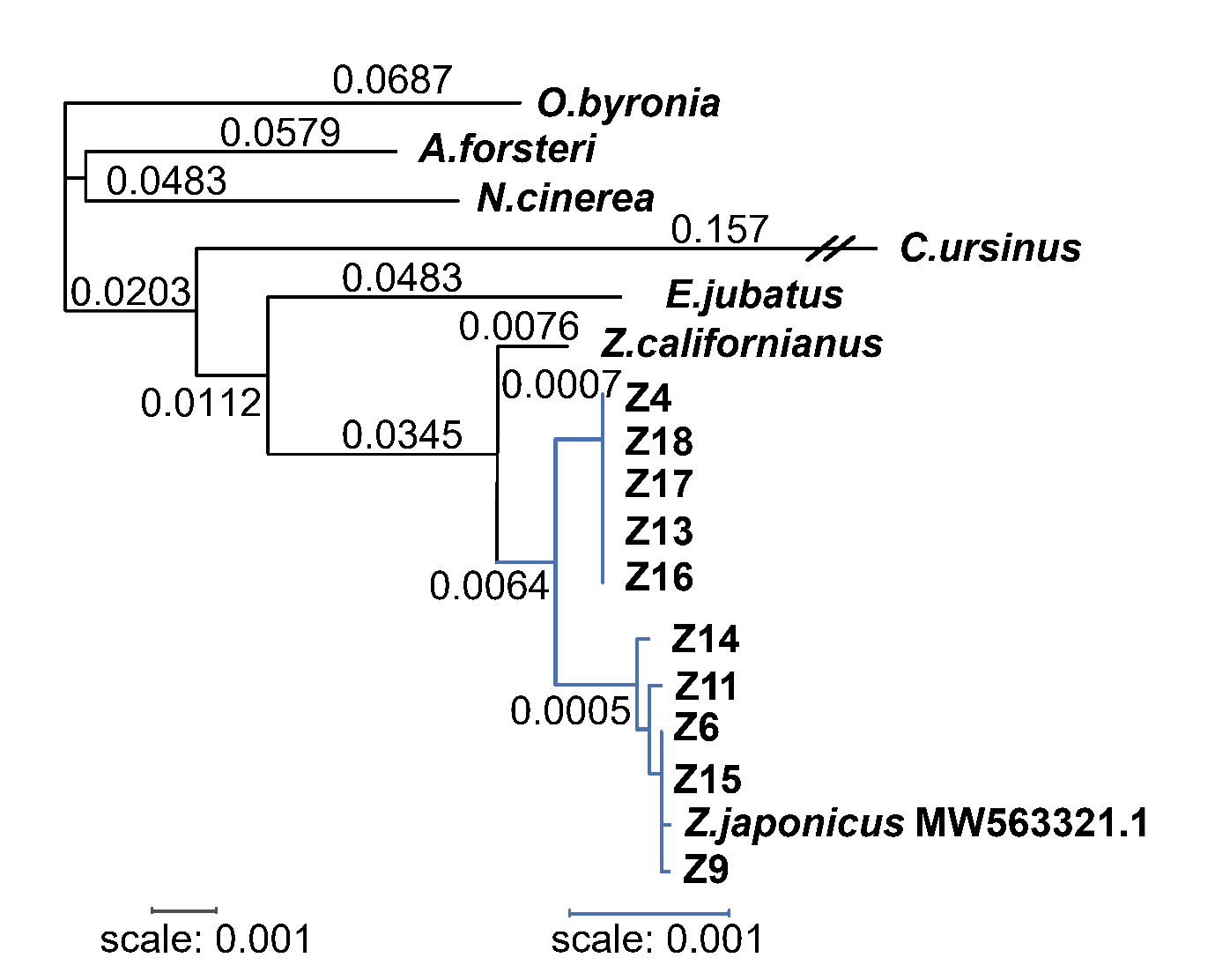
 **Fig. S6. Phylogenetic tree of the mitogenomes of Dokdo sea lion and related species**

This phylogenetic illustrates the evolutionary relationships among *Z. japonicus* individuals (Z4, Z6, Z9, Z11, Z13-18 and previously published MW563321.1), now extinct, and its closest relatives based on mitochondrial DNA sequences. The length of each branch corresponds to the genetic distance between the species. Each blue branch represents a species within the *Z. japonicus* and black branches represents genetic distance with related species. The tree shows that *Z. japonicus*, *Z. californianus* are closely related, sharing a common ancestor of *E. jubatus*.

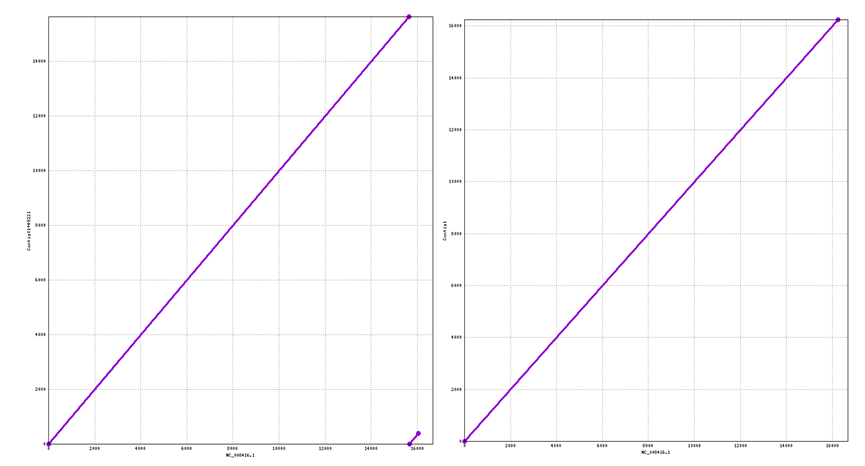
 **Fig. S7. Comparison of mitochondrial genomes between *Z. japonicus* and *Z. californianus* using mummer (ver. 4.0.0rc1)**

It showed mitochondrial genome alignment between *Z. californianus* and *Z. japonicus* assembly. The left alignment was Pre-gap filling alignment and the right one was post-gap filling. The completed mitochondrial genome assembly of the California sea lion along the x-axis, and the mitochondrial genome contigs of *Z. japonicus* on the y-axis. This alignment demonstrates the enhanced contiguity and sequence integrity achieved after gap filling, indicative of a more accurate and comprehensive mitochondrial genome assembly for both species.

Supplementary Tables

Table S1. Statistics of the PALEOMIX pipeline mapping to California sea lion (*Z. californianus*) reference genome

| Description | Statistics |
| --- | --- |
| SE, PE, or * (for both) |  |
| Total number of pairs | 27,429,982,138 |
| Total number of reads | 2,128,601 |
| Fraction of PE mate 1 reads trashed | 7.76e^-05^ |
| Total number of reads | 2,128,601 |
| Fraction of PE mate 2 reads trashed | 7.76e^-05^ |
| Total number of pairs collapsed into one read | 11,295,109,116 |
| Fraction of PE pairs collapsed into one read | 0.41 |
| Total number of retained reads | 43,560,597,958 |
| Total number of NTs in retained reads | 6,925,953,587,955 |
| Average number of NTs in retained reads | 158.9958337 |
| Total number of hits (prior to PCR duplicate filtering) | 33,148,490 |
| Total number of hits vs. total number of reads retained | 0.000760974 |
| Fraction of hits that were PCR duplicates | 0.10 |
| Total number of hits (excluding any PCR duplicates) | 29,887,682 |
| Total number of unique hits vs. total number of reads retained | 0.000686117 |
| Estimated coverage from unique hits | 1.73 |
| Average number of aligned bases per unique hit | 139.66 |

Table S2. Gene flow between Steller sea lion (*E. jubatus*) and *Zalophus* species with a form of *f4*(A,B;C,D). A positive Z-score > 3 indicates significant gene flow from B to D. Red text indicates statistically significant gene flow. The species abbreviations are as follows: *C. urs* (*C. ursinus*); *E. jub* (*E. jubatus*); *Z. jap* (*Z. japonicus*); *Z. cal* (*Z. californianus*); *Z. wol* (*Z. wollebaeki*).

| A | B | C | D | *f4* stat | Z-score | BABA | ABBA | est |
| --- | --- | --- | --- | --- | --- | --- | --- | --- |
| *C. urs* | *Z. jap* | *E. jub* | *Z. cal* | 0.724 | 96.416 | 27,535 | 4,407 | 1,581,963 |
| *C. urs* | *Z. jap* | *E. jub* | *Z. wol* | 0.7227 | 94.502 | 27,291 | 4,391 | 1,581,963 |
| *C. urs* | *Z. cal* | *E. jub* | *Z. wol* | 0.6945 | 100.000 | 31,888 | 5,748 | 1,581,963 |
| *C. urs* | *Z. cal* | *E. jub* | *Z. jap* | 0.6173 | 75.998 | 27,535 | 6,514 | 1,581,963 |
| *C. urs* | *Z. wol* | *E. jub* | *Z. cal* | 0.6974 | 100.000 | 31,888 | 5,683 | 1,581,963 |
| *C. urs* | *Z. wol* | *E. jub* | *Z. jap* | 0.6185 | 75.346 | 27,291 | 6,433 | 1,581,963 |

Table S3. *f4*-statistics of *Otariidae* species with a form of f4(A,B;C,D). A positive Z-score > 3 indicates significant gene flow from B to D, while a negative Z-score < -3 indicates significant gene flow from B to C. Red text indicates statistically significant gene flow. The species abbreviations are as follows: *C. urs* (*C. ursinus*); *E. jub* (*E. jubatus*); *Z. jap* (*Z. japonicus*); *Z. cal* (*Z. californianus*); *Z. wol* (*Z. wollebaeki*).

| A | B | C | D | f4 stat | Z-score | BABA | ABBA | SNPs |
| --- | --- | --- | --- | --- | --- | --- | --- | --- |
| *C. urs* | *E. jub* | *Z. jap* | *Z.cal* | 0.1928 | 31.324 | 6,514 | 4,407 | 1,581,963 |
| *C. urs* | *E. jub* | *Z. jap* | *Z.wol* | 0.1884 | 28.809 | 6,433 | 4,391 | 1,581,963 |
| *C. urs* | *E. jub* | *Z. wol* | *Z.cal* | 0.0057 | 1.993 | 5,748 | 5,683 | 1,581,963 |
| *E. jub* | *Z.jap* | *Z.cal* | *Z. wol* | -0.0125 | -3.266 | 6,425 | 6,588 | 1,581,963 |
| *E. jub* | *Z.wol* | *Z. jap* | *Z.cal* | 0.2012 | 34.596 | 9,665 | 6,425 | 1,581,963 |
| *E. jub* | *Z.cal* | *Z.wol* | *Z. jap* | -0.1892 | -35.743 | 6,588 | 9,665 | 1,581,963 |

Table S4. Genetic admixture of *Zalophus* species compared to their related species. A Z-score < -3 indicates that the ancestral lineage of “T” has been admixed by lineages A and B. The species abbreviations are as follows: *C. urs* (*C. ursinus*); *E. jub* (*E. jubatus*); *Z. jap* (*Z. japonicus*); *Z. cal* (*Z. californianus*); *Z. wol* (*Z. wollebaeki*).

| A | B | Target | f_3 | std. err | Z | SNPs |
| --- | --- | --- | --- | --- | --- | --- |
| *C. urs* | *Z. jap* | *Z. cal* | 0.069815 | 0.002037 | 34.274 | 396,284 |
| *C. urs* | *Z. wol* | *Z. cal* | -0.00956 | 0.00128 | -7.474 | 392,924 |
| *C. urs* | *Z. jap* | *Z. wol* | -0.02296 | 0.004741 | -4.842 | 338,734 |
| *C. urs* | *Z. cal* | *Z. wol* | -0.09926 | 0.003388 | -29.293 | 392,924 |
| *C. urs* | *Z. cal* | *Z. jap* | 0.140503 | 0.012199 | 11.518 | 396,284 |
| *C. urs* | *Z. wol* | *Z. jap* | 0.147069 | 0.012253 | 12.002 | 338,734 |
| *E. jub* | *Z. jap* | *Z. cal* | 0.037144 | 0.001757 | 21.146 | 264,510 |
| *E. jub* | *Z. wol* | *Z. cal* | -0.01058 | 0.001262 | -8.383 | 259,703 |
| *E. jub* | *Z. jap* | *Z. wol* | -0.05208 | 0.004182 | -12.453 | 210,702 |
| *E. jub* | *Z. cal* | *Z. wol* | -0.09832 | 0.003395 | -28.957 | 259,703 |
| *E. jub* | *Z. cal* | *Z. jap* | 0.201034 | 0.0141 | 14.258 | 264,510 |
| *E. jub* | *Z. wol* | *Z. jap* | 0.205712 | 0.014179 | 14.509 | 210,702 |
| *Z. wol* | *Z. cal* | *Z. jap* | 0.294155 | 0.016306 | 18.039 | 187,733 |
| *Z. jap* | *Z. cal* | *Z. wol* | -0.096 | 0.003447 | -27.852 | 187,733 |
| *Z. wol* | *Z. jap* | *Z. cal* | -0.0131 | 0.001251 | -10.47 | 187,733 |

Table S5. Primers to filling the mito-genome gap of Dokdo sea lion (Z6 individual) and melting temperature (Tm) used

| Forward primer (FWD), 5’- 3’ | Reverse primer (REV), 5’ – 3’ | Tm | | PCR-fragment length, bp |
| --- | --- | --- | --- | --- |
|  |  | FWD | REV |  |
| GGGGACTGGTATCACTCAGC | ACGGAAGGGCTAGGACCA | 59.534 | 59.552 | 408 |

Table S6. Sanger sequencing reads to fill mito-genome gap of Z6 individual

>CalZ-Gap_filling_F

CATGGTCTGGACAGTCAATAACTTGTAGCTGGACTTAATTATTATCATTTACCAGCATCATACAACCATGAGGCGCATTTTAGTCAATGGTCGCAGGACATACACGTATACGCACACACGTATACACGTATACACGTATACACGTATACGCATACACGCATACACGTATACACGTATACACGTATACACGTATACGCGCACACGCATACACGTATACACGTATACACGTATACACGTATACGCGCATACGCATACACGTATACACGTATACACGTATACACGTATACACGCATACACATATACACATATACACATATACGCATATACACATATACGCATATACGCATATACGCATATACACATATACGCATATACGCGTATACGCATATACGTATATACGTATACACGTATACACGTATACACGTATACACGTATACACGTATACATGTACATGTGTACGTGAATGCCTGACACTTCGAAAACTAGAAACATCTCCCCTTATCCCCTGAAACTCCGAAAATTATAAACATATGAACTGCTGCACTGCTGAACTGCAGAAATGCAGATATGTAAATGCGAAACAGATGAGCCGAGGAGCCGCAGAGCAGTTACGCATTACCAAACTAGTATACATGTCTACATATATATATATATCGATTTATACTTATCTCTATCTATATATCTATCTATTTCGCTTCTACTCTCTCATATTATCATTCTACTATTCTACCATCTTACACTCAATAGAGCAGTGACTCAGTGTAGCTGAGTCACTGTCACAGCTGAGGCACTGAAGCTGTGTAGCTGAGTCGCGAGACTCGGAACTGCGAGATTTGACGTAGGACATGCGTATT

>CalZ-Gap_filling_R

CCGGGTATGGGAGTCTCGTGACTCATCTAGGCATTTTCAGTGCCTTGCTTTGAGTTTAGTTAAGCTACATTAACTGGTTGATTTTATGGGGGTGGTTGGTGGGAATAAAAGTGGAAGGTTAATAGGGGGGAGTTAGAGGCGGGAAATGTATATCTATAGATAGACTTAAGATTTAAGTATTATGTTTGTGTTTTGTGACCTGTTGACTGGGTTGGTTAAGTTAAATTTGGTTGCAGATCCTGGCTTCATCTTTGTTGCATTTATCTAGTTCTGTTTTTGGGGTTTGGCAGGACAAAAGTGGATATCTGTATACTTTTTGAAGTTACGGGGGGTAAGGGGGGTTTGTCTTAGTTAACTTAATGTCTGTCATTTAAACTAACATGAATATGTCTACGTGTATACGTGTATACGTGTATACGTGTATACGTGTATACGTGTATACGTGTATACGTGTATACGTGTATACGTGTATACGTGTATACGTGTATACGTGTATACGTGTATACGTGTATACGTGTATACGTGTATACGTGTATACGTGTATGCGTGTACGCGTATACGTGTATACGTGTATACGTGTATACGTGTATACGTGTACGCGTATACGTGTATACGTGTATACGTGTATACGTGTATGCGTGTACGCGTATACGTGTATACGTGTATACGTGTATACGTGTATACGTGTATGCGTATACGCGTATACGTGTATACGTGTGTGCGTGTATGTGTATATCCTGCTATCATTGTATAATATTCTACTCCTGGAATTATGAAGTTAAGAAACAAAAAAACTAAAATCCAGTAACTCAGTATCTCAGTGTCTCAGGACCTCAAGAACCCGAAGGTGAGGGGAACCGGGGCCCAGGGCCCCAGGGACCGAAG
